# Genomic epidemiology supports multiple introductions and cryptic transmission of Zika virus in Colombia

**DOI:** 10.1101/454777

**Authors:** Allison Black, Louise H. Moncla, Katherine Laiton-Donato, Barney Potter, Lissethe Pardo, Angelica Rico, Catalina Tovar, Diana P. Rojas, Ira M. Longini, M. Elizabeth Halloran, Dioselina Peláez-Carvajal, Juan D. Ramírez, Marcela Mercado-Reyes, Trevor Bedford

## Abstract

**Background:** Colombia was the second most affected country during the American Zika virus (ZIKV) epidemic, with over 109,000 reported cases. Despite the scale of the outbreak, limited genomic sequence data were available from Colombia. We sought to sequence additional samples and use genomic epidemiology to describe ZIKV dynamics in Colombia.

**Methods:** We sequenced ZIKV genomes directly from clinical diagnostic specimens and infected *Aedes aegypti* samples selected to cover the temporal and geographic breadth of the Colombian outbreak. We performed phylogeographic analysis of these genomes, along with other publicly-available ZIKV genomes from the Americas, to estimate the frequency and timing of ZIKV introductions to Colombia.

**Results:** We attempted PCR amplification on 184 samples; 19 samples amplified sufficiently to perform sequencing. Of these, 8 samples yielded sequences with at least 50% coverage. Our phylogeographic reconstruction indicates two separate introductions of ZIKV to Colombia, one of which was previously unrecognized. We find that ZIKV was first introduced to Colombia in February 2015 (95%CI: Jan 2015 - Apr 2015), corresponding to 5 to 8 months of cryptic ZIKV transmission prior to confirmation in September 2015. Despite the presence of multiple introductions, we find that the majority of Colombian ZIKV diversity descends from a single introduction. We find evidence for movement of ZIKV from Colombia into bordering countries, including Peru, Ecuador, Panama, and Venezuela.

**Conclusions:** Similarly to genomic epidemiologic studies of ZIKV dynamics in other countries, we find that ZIKV circulated cryptically in Colombia. This more accurate dating of when ZIKV was circulating refines our definition of the population at risk. Additionally, our finding that the majority of ZIKV transmission within Colombia was attributable to transmission between individuals, rather than repeated travel-related importations, indicates that improved detection and control might have succeeded in limiting the scale of the outbreak within Colombia.

## Background

In recent years, countries across the Americas have experienced the emergence and endemic circulation of various mosquito-borne viruses, making this a critical area for public health surveillance and epidemiologic research. Zika virus (ZIKV) caused a particularly widespread epidemic, with over 800,000 suspected or confirmed cases reported [1]. Given estimated seroprevalence rates between 36% and 76% [2–5], the true number of ZIKV infections in the Americas is likely much higher. With neither a vaccine nor ZIKV-specific treatments available, understanding the epidemiology of ZIKV is our primary tool for controlling disease spread [6]. However, because many infections are asymptomatic [2], the analysis of surveillance data alone yields inaccurate estimates of when ZIKV arrived in a country [7, 8]. In such cases, introduction timing and transmission dynamics post-introduction are better inferred from genomic epidemiological studies, which use joint analysis of viral genome sequences and epidemiologic case data. Indeed, such studies have defined our understanding of when ZIKV arrived in Brazil [7], described general patterns of spread from Brazil into other countries in the Americas [7, 9, 10], and been used to investigate the extent of endemic transmission occurring post-introduction [8]. Genomic epidemiological studies of the spread of ZIKV in the Americas have aided our understanding of the epidemic [7–19], but generally, ZIKV pathogen sequencing has remained a challenge for the public health community [20].

Colombia has a population of approximately 48 million people. In addition, Colombia has *Aedes aegypti* and *Ae.albopictus* mosquitoes, which are commonly found at elevations below 2000m above sea level [21]. Public health surveillance for arboviral diseases, along with other notifiable conditions, is performed by the Instituto Nacional de Salud de Colombia (INS) [21]. While suspected cases from other municipalities were reported earlier [22], the INS first confirmed ZIKV circulation in mid-September 2015, in the Turbaco municipality on the Caribbean coast. ZIKV spread throughout the country, appearing in areas infested with *Ae.aegypti* that experience endemic dengue transmission and previous circulation of chikungunya virus [23]. Over the entire epidemic Colombia reported 109,265 cases of Zika virus disease [24], making it the second most ZIKV-affected country in the Americas after Brazil. The extent of the epidemic led the INS to start active surveillance for congenital Zika syndrome [25] as well as other neurological syndromes associated with ZIKV infection [26]. While the INS determined that epidemic ZIKV transmission ended in July 2016, they continue to perform surveillance for endemic transmission.

Despite numerous reported cases, only 12 whole ZIKV genomes from Colombian clinical samples were publicly available. These sequences included 1 sample from Barranquilla, Atlántico department, 4 samples from Santander department, and 7 sequences for which departmental or municipal information was unspecified. We sequenced an additional 8 samples from ZIKV-positive human clinical and *Ae.aegypti* specimens, sampled from previously unrepresented Colombian departments. We describe here the first detailed phylogeographic analysis of Colombian ZIKV to estimate when, and how frequently, ZIKV was introduced into Colombia.

## Methods

### INS sample selection and processing

The INS National Virological Reference Laboratory collected diagnostic specimens from over 32,000 suspected ZIKV cases over the course of the epidemic. Of these, roughly 800 serum specimens were positive for ZIKV by real time RT-PCR (rRT-PCR) and had cycle threshold (Ct) values less than 30. From this set we selected 176 serum specimens that were ZIKV-positive, but negative for dengue and chikungunya viruses, as per results from the Trioplex rRT-PCR assay [27]. Specimens were selected such that each Colombian department was represented over the entire time period in which specimens were submitted. We extracted RNA using the MagNA Pure 96 system (Roche Molecular Diagnostics, Pleasanton, CA, USA) according to manufacturers instructions. An extraction negative was used for each plate; positive controls were eschewed given the risk of cross-contaminating low titer clinical samples [20]. We attempted reverse transcription and PCR-amplification of ZIKV using the two-step multiplex PCR protocol developed by Quick and colleagues on all 176 extracted samples [20]. Briefly, cDNA was generated using random hexamer priming and the Protoscript II First Strand cDNA Synthesis Kit (New England Biolabs, Ipswich, MA, USA). We amplified cDNA using the ZikaAsian V1 ZIKV-specific primer scheme [20], which amplifies 400bp long overlapping amplicons across the ZIKV genome, over 35 cycles of PCR. Amplicons were purified using 1x AMPure XP beads (Beckman Coulter, Brea, CA, USA) and quantified using with the Qubit dsDNA High Sensitivity assay on the Qubit 3.0 instrument (Life Technologies, Carlsbad, CA, USA). Due to long storage periods and variable storage temperature, the vast majority of samples were too degraded to amplify. Of the 176 processed samples, only 15 amplified sufficiently to perform sequencing.

### UR sample selection and processing

Universidad del Rosario (UR) collected and performed diagnostic testing on 23 human clinical samples from different geographic regions, and 38 *Ae.aegypti* samples from the Cordoba department of Colombia. RNA was extracted using the RNeasy kit (Qiagen, Hilden, Germany) and a single TaqMan assay (Applied Biosystems, Foster City, CA, USA) directed to ZIKV was employed [28] to confirm ZIKV presence. Approximately 60% of samples (14 clinical samples and 23 *Ae.aegypti* samples) were found to be ZIKV-positive by rRT-PCR. From these, we attempted amplification on 8 samples with sufficiently high viral copy numbers that they were likely to amplify. Amplification, purification, and quantification of ZIKV amplicons from UR samples were performed as described above. Of the eight samples that we performed PCR on, four samples amplified sufficiently to conduct sequencing; three samples were from human clinical specimens and one was from an *Ae.aegypti* sample.

### Sequencing protocol

We sequenced amplicons from 4 UR and 15 INS samples using the Oxford Nanopore MinION (Oxford Nanopore Technologies, Oxford, UK) according to the protocol described in Quick et al [20]. Amplicons were barcoded using the Native Barcoding Kit EXP-NBD103 (Oxford Nanopore Technologies, Oxford, UK) and pooled in equimolar fashion. Sequencing libraries were prepared using the 1D Genomic DNA Sequencing kit SQK-LSK108 (Oxford Nanopore Technologies, Oxford, UK). We used AMPure XP beads (Beckman Coulter, Brea, CA, USA) for all purification steps performed as part of library preparation. Prepared libraries were sequenced on R9.4 flowcells (Oxford Nanopore Technologies, Oxford, UK) at the INS in Bogotá and at the Fred Hutchinson Cancer Research Center in Seattle.

### Bioinformatic processing

Raw signal level data from the MinION were basecalled using Albacore version 2.0.2 (Oxford Nanopore Technologies, Oxford, UK) and demultiplexed using Porechop version 0.2.3 seqan2.1.1 github.com/rrwick/Porechop. Primer binding sites were trimmed from reads using custom scripts, and trimmed reads were mapped to Zika reference strain H/PF/2013 (GenBank Accession KJ776791) using BWA v0.7.17 [29]. We used Nanopolish version 0.9.0 github.com/jts/nanopolish to determine single nucleotide variants from the event-level data, and used custom scripts to extract consensus genomes given the variant calls and the reference sequence. Coverage depth of at least 20x was required to call a SNP; sites with insufficient coverage were masked with N, denoting that the exact nucleotide at that site is unknown. After bioinformatic assembly, 8 samples produced sufficiently complete genomes to be informative for phylogenetic analysis.

### Dataset curation

All publicly available Asian lineage ZIKV genomes and their associated metadata were downloaded from ViPR [30] and NCBI GenBank. The full download contained both published and unpublished sequences; we sought written permission from submitting authors to include sequences that had not previously been published on. Any sequences for which we did not receive approval were removed. Additionally, we excluded sequences from the analysis if any of the following conditions were met: the sequence had ambiguous base calls at half or more sites in the alignment, the sequence was from a cultured clone for which a sequence from the original isolate was available, the sequence was sampled from countries outside the Americas or Oceania, or geographical sampling information was unknown. Finally, we also excluded viruses whose estimated clock rate was more than 4 times the interquartile distance of clock rates across all sequences in the analysis. Sequences that deviate this greatly from the average evolutionary rate either have far too many or too few mutations than expected given the date they were sampled. This deviation usually occurs if the given sampling date is incorrect, or if the sequence has been affected by contamination, lab adaptation, or sequencing error. After curation, the final dataset consisted of 360 sequences; 352 publicly available ZIKV full genomes from the the Americas (including Colombia) and Oceania, and the 8 Colombian genomes from the present study.

### Phylogeographic analysis

Data were cleaned and canonicalized using Nextstrain Fauna github.com/nextstrain/fauna, a databasing tool that enforces a schema for organizing sequence data and sample metadata, thereby creating datasets compatible with the Nextstrain Augur analytic pipeline github.com/nextstrain/augur and the Nextstrain Auspice visualization platform github.com/nextstrain/auspice. A full description of the Nextstrain pipelines can be found in Hadfield et al [31].Briefly, Nextstrain Augur performs a multiple sequence alignment with MAFFT [32], which is then trimmed to the reference sequence. A maximum likelihood phylogeny is inferred using IQ-TREE [33]. Augur then uses TreeTime [34] to estimate a molecular clock; rates inferred by TreeTime are comparable to BEAST [34], a program that infers temporally-resolved phylogenies in a Bayesian framework [35]. Given the inferred molecular clock, TreeTime then creates a temporally-resolved phylogeny, infers sequence states at internal nodes, and estimates the geographic migration history across the tree. These data are exported as JSON files that can be interactively visualized on the web using Nextstrain Auspice.

### Rarefaction analysis

Briefly, we generated a series of datasets in which ZIKV genomes from either Colombia or Mexico were subsampled. For each value from 1 to *n*, where *n* is the total available sequences for the country of interest, we generated 30 subsampled sets in which *n* sequences were randomly sampled without replacement. Each subsampled set was then combined with all ZIKV genomes from other countries included in the phylogeographic analysis described above, and the phylogeographic inference was rerun. For the Colombian rarefaction analysis, we have *n* = 20 high quality whole genomes, so we inferred 20 · 30 = 600 phylogeographically labelled trees. For the Mexican rarefaction analysis, we have *n* = 51 high quality whole genomes in total, and inferred 51 · 30 = 1530 phylogeographically labelled trees. For each labelled tree, we used custom scripts to traverse the resulting phylogeny and count the number of ZIKV introductions into the country of interest, and plotted the inferred introduction count as a function of the number of sequences sampled.

## Results

### Sequencing and sampling characteristics of reported ZIKV genomes

In total, we attempted to amplify ZIKV nucleic acid from 184 samples collected by the Instituto Nacional de Salud de Colombia (INS) and Universidad del Rosario (UR). Given the low viral titers associated with most ZIKV infections, as well as long storage times, most samples did not amplify well. We attempted sequencing on 19 samples that amplified sufficiently to generate sequencing libraries (Table 1). Sequencing efforts yielded eight ZIKV sequences with at least 50% coverage across the genome with unambiguous base calls (Table 1). Seven of these viruses came from humans; one virus came from an *Ae.aegypti* pool. Three sequences came from samples collected from infected individuals in Cali, department of Valle del Cauca, two sequences came from Montería, department of Córdoba, and one sequence each came from Ibagué, department of Tolima, Belén de Umbría, department of Risaralda, and Pitalito, department of Huila (Figure 1). Colombian viruses are sampled across the period of peak ZIKV incidence in Colombia (Figure 2).

**Table 1.**
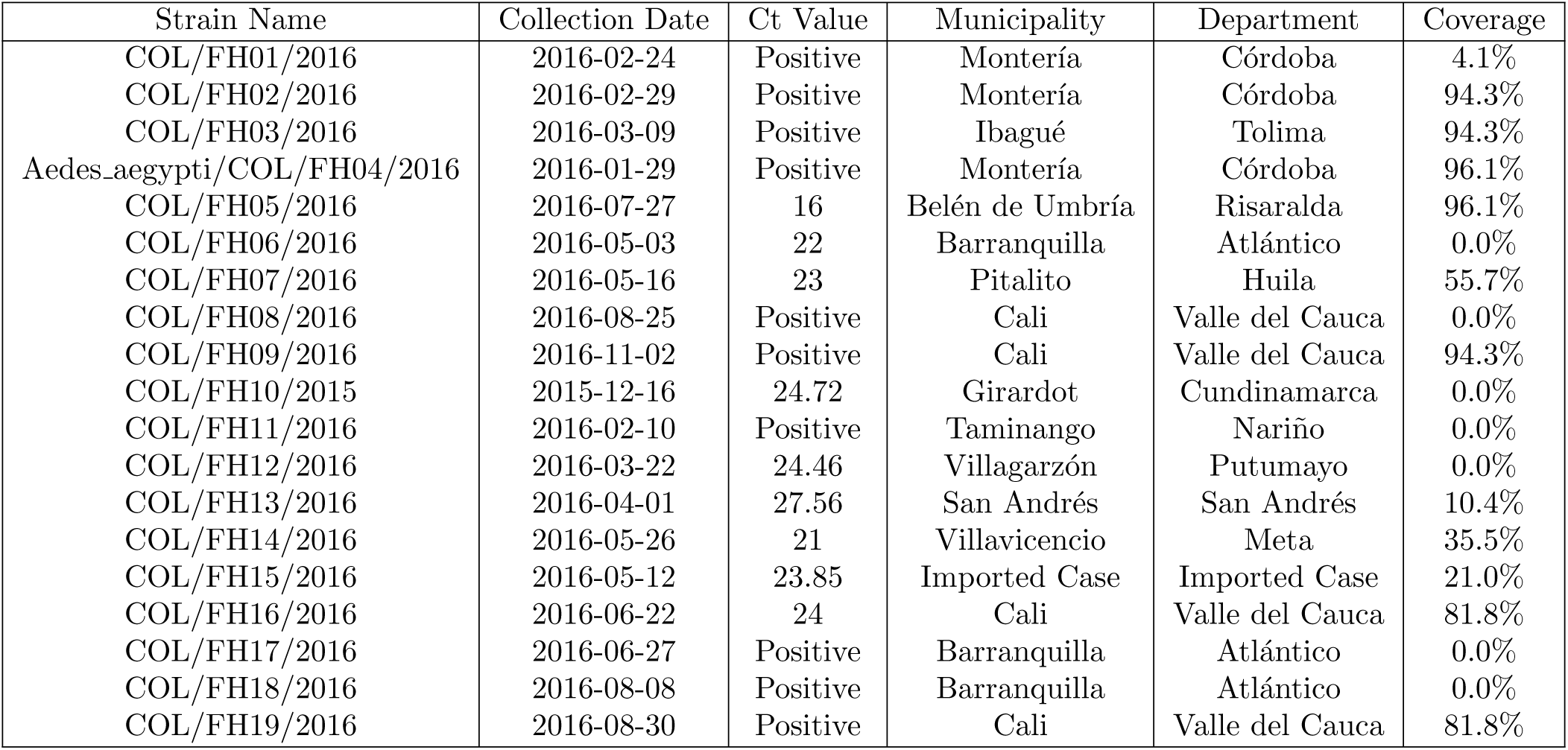
Metadata and percent of genome that was able to be called unambiguously for 19 samples that amplified sufficiently to attempt sequencing. Not all samples had annotated Ct values; rather, in some cases, samples were only specified to be positive or negative for ZIKV.

| Strain Name | Collection Date | Ct Value | Municipality | Department | Coverage |
| --- | --- | --- | --- | --- | --- |
| COL/FH01/2016 | 2016-02-24 | Positive | Montería | Córdoba | 4.1% |
| COL/FH02/2016 | 2016-02-29 | Positive | Montería | Córdoba | 94.3% |
| COL/FH03/2016 | 2016-03-09 | Positive | Ibagué | Tolima | 94.3% |
| Aedes.aegypti/COL/FH04/2016 | 2016-01-29 | Positive | Montería | Córdoba | 96.1% |
| COL/FH05/2016 | 2016-07-27 | 16 | Belén de Umbría | Risaralda | 96.1% |
| COL/FH06/2016 | 2016-05-03 | 22 | Barranquilla | Atlántico | 0.0% |
| COL/FH07/2016 | 2016-05-16 | 23 | Pitalito | Huila | 55.7% |
| COL/FH08/2016 | 2016-08-25 | Positive | Cali | Valle del Cauca | 0.0% |
| COL/FH09/2016 | 2016-11-02 | Positive | Cali | Valle del Cauca | 94.3% |
| COL/FH10/2015 | 2015-12-16 | 24.72 | Girardot | Cundinamarca | 0.0% |
| COL/FH11/2016 | 2016-02-10 | Positive | Taminango | Nariño | 0.0% |
| COL/FH12/2016 | 2016-03-22 | 24.46 | Villagarzón | Putumayo | 0.0% |
| COL/FH13/2016 | 2016-04-01 | 27.56 | San Andrés | San Andrés | 10.4% |
| COL/FH14/2016 | 2016-05-26 | 21 | Villavicencio | Meta | 35.5% |
| COL/FH15/2016 | 2016-05-12 | 23.85 | Imported Case | Imported Case | 21.0% |
| COL/FH16/2016 | 2016-06-22 | 24 | Cali | Valle del Cauca | 81.8% |
| COL/FH17/2016 | 2016-06-27 | Positive | Barranquilla | Atlántico | 0.0% |
| COL/FH18/2016 | 2016-08-08 | Positive | Barranquilla | Atlántico | 0.0% |
| COL/FH19/2016 | 2016-08-30 | Positive | Cali | Valle del Cauca | 81.8% |

**Figure 1.**
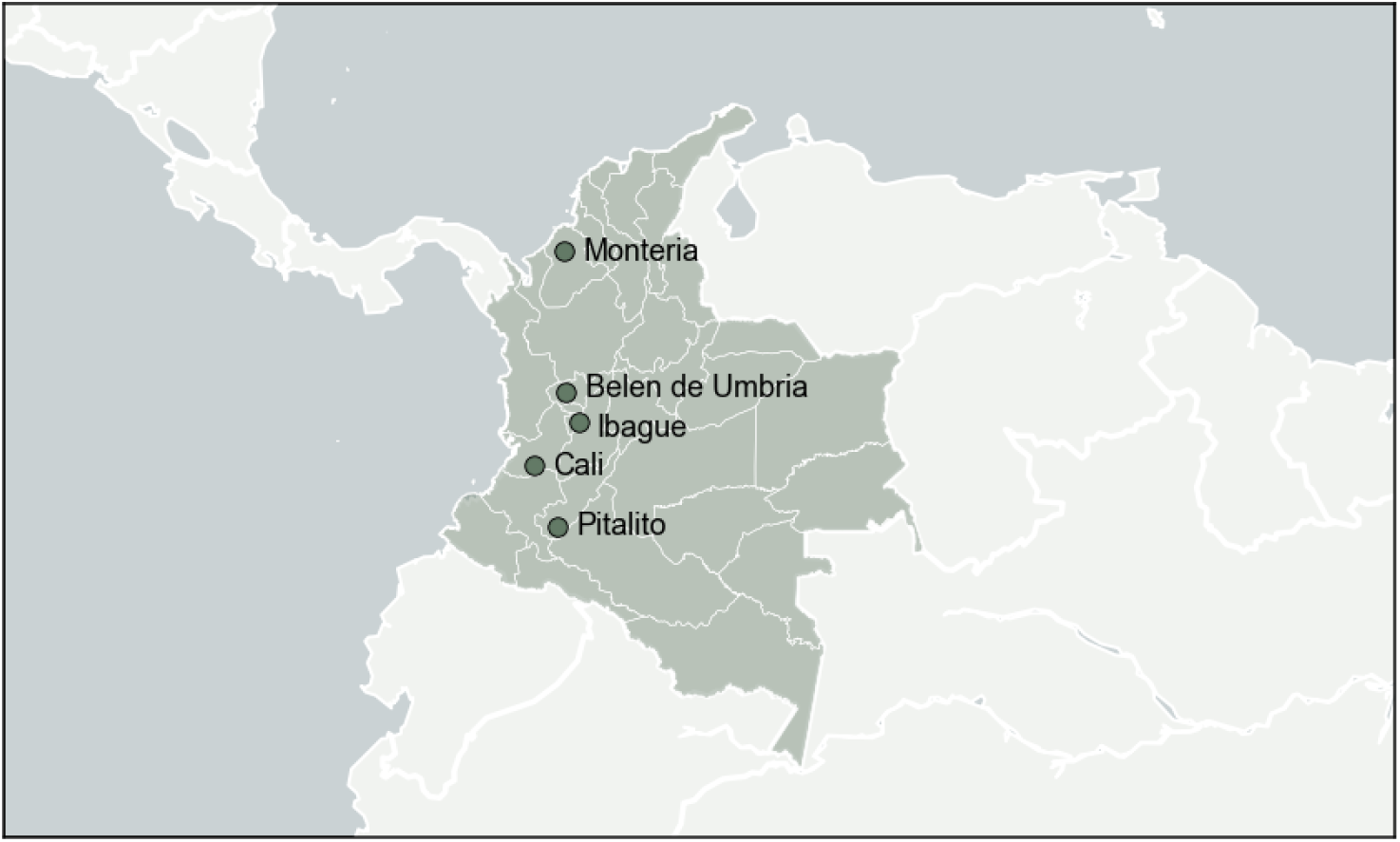
Geographic sampling locations for the eight Colombian genomes with at least 50% unambiguous genome coverage. Colombia is highlighted with departmental boundaries shown. Three genomes come from Cali and and two genomes come from Montería. All other cities have one genome each. The land-sea mask, coastline, lake, river and political boundary data are extracted from datasets provided by Generic Mapping Tools (GMT) licensed under GNU General Public License.

**Figure 2.**
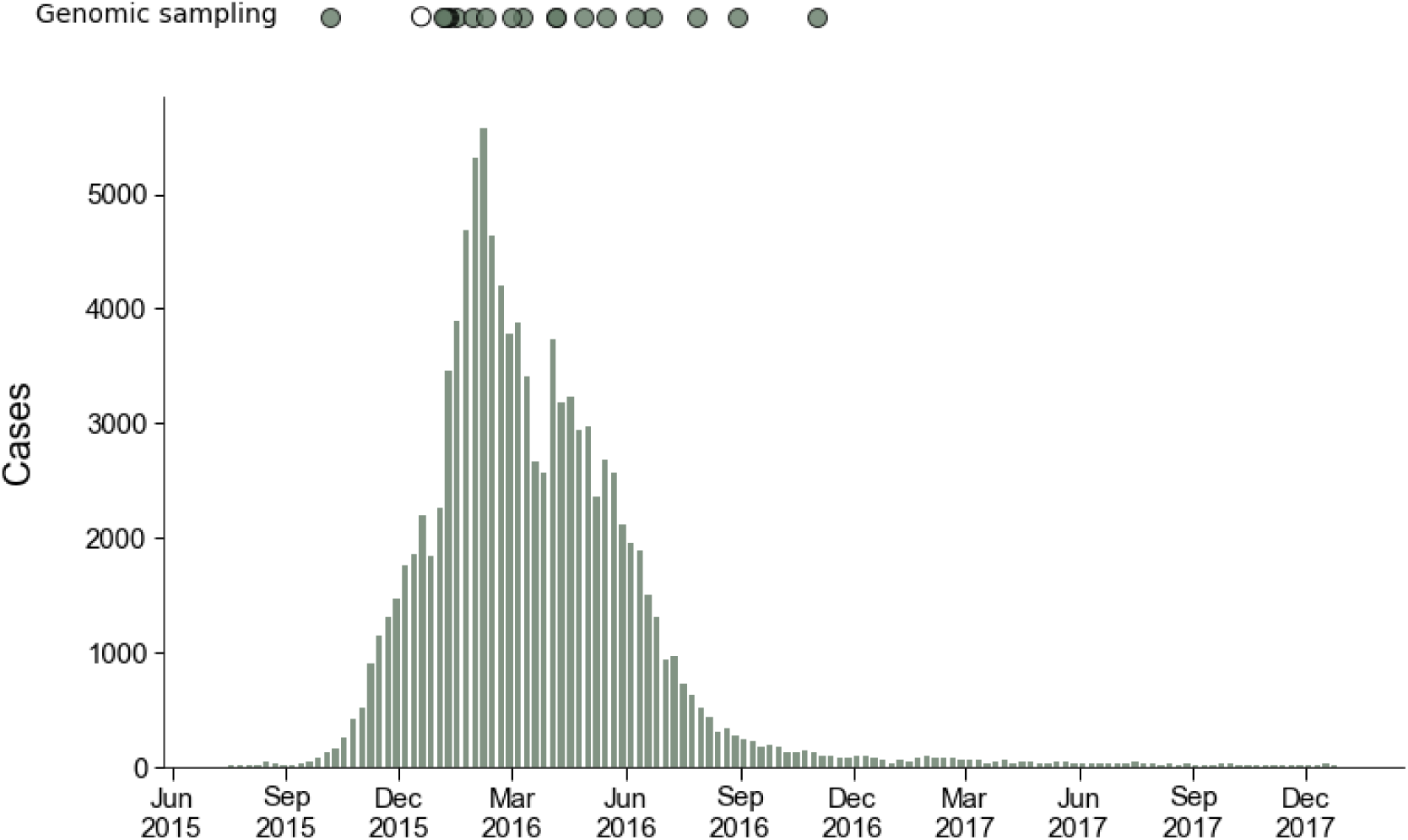
Numbers of recorded cases weekly from the beginning of the Colombian epidemic to the end of 2017. Sampling dates for 20 Colombian genomes, including the eight sequenced as part of this study, are indicated by circles above the epidemiologic curve. One sample lacked information about sampling date, and sampling date was inferred. This sample is indicated as an unfilled circle.

### General patterns of ZIKV transmission in the Americas

We conducted a maximum likelihood phylogeographic analysis of 360 Asian lineage ZIKV genomes; 18 sequences are sampled from Oceania, and 342 sequences are sampled from the Americas. We estimate the ZIKV evolutionary rate to be 8.65 × 10^−4^ substitutions per site per year, in agreement with other estimates of the ZIKV molecular clock [7, 10, 11, 36] made with BEAST [35], a Bayesian method for estimating rates of evolution and inferring temporally-resolved phylogenies. Consistent with other studies, we find that ZIKV moved from Oceania to the Americas, and that the American epidemic descends from a single introduction into Brazil (Figure 3A). We estimate that this introduction occurred in late September 2013 (95%CI: August 2013 - March 2014), inline with Faria et al’s [11] initial estimate of introduction to Brazil between May and December 2013, and updated estimate of introduction between October 2013 and April 2014 [7]. We also confirm findings from previous studies [7, 10] that ZIKV circulated in Brazil for approximately one year before moving into other South American countries to the north, including Colombia, Venezuela, Suriname, and French Guiana. Movement of ZIKV into Central America occurs around late-2014 while movement into the Caribbean occurs slightly later, around mid-2015 (Figure 3A).

**Figure 3.**
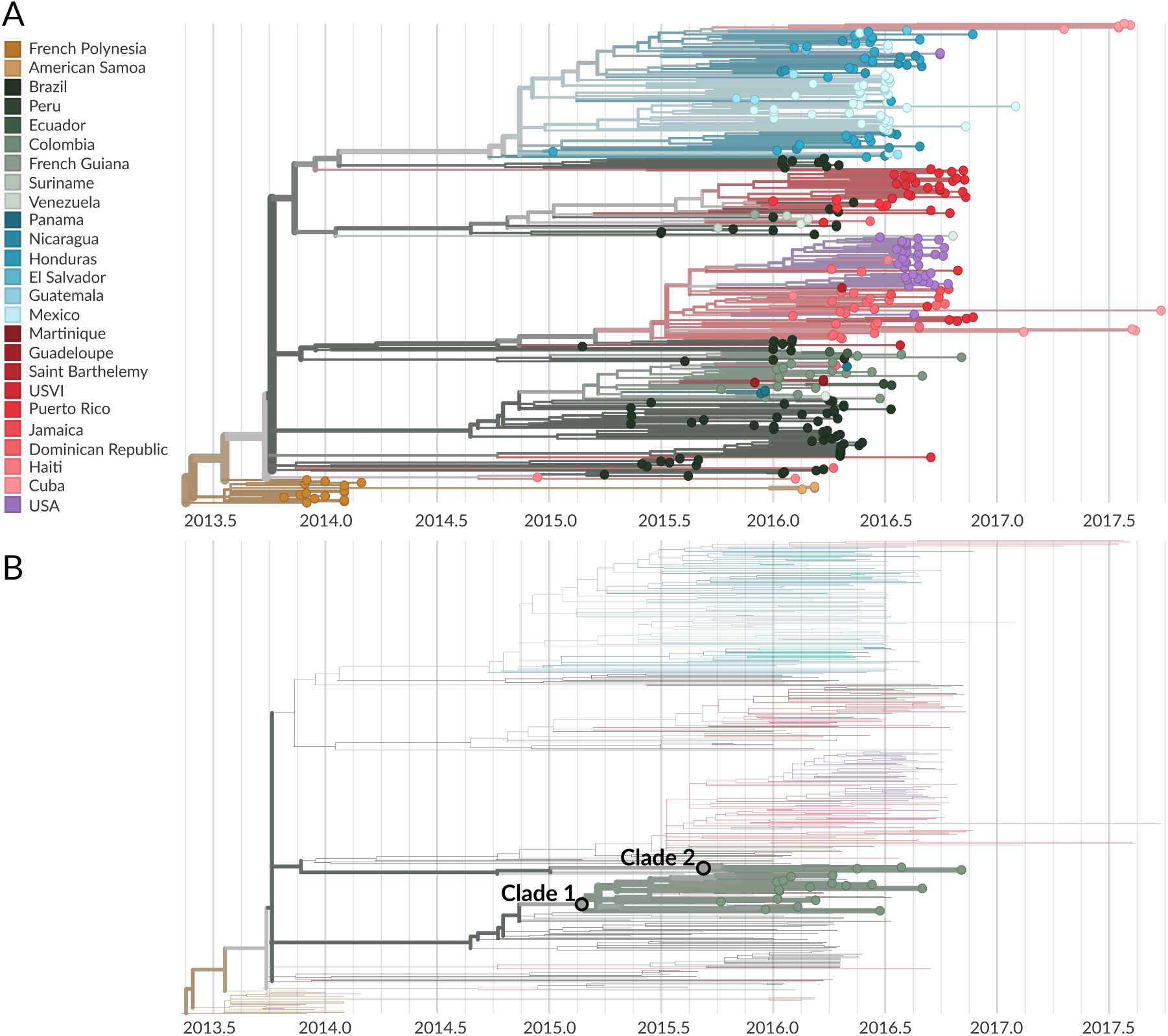
Phylogeographic analysis of 360 publicly available ZIKV genomes. A) Temporally-resolved maximum likelihood phylogeny of 360 ZIKV genomes from the Americas and Oceania. Tip colors indicate known country of sampling and branch colors indicate geographic migration history inferred under the phylogeographic model. The full tree can be explored interactively at nextstrain.org/community/blab/zika-colombia. B) The same phylogeny as in panel A, but filtered to highlight only genomes sampled from Colombia. The phylogeny shows two separate introductions of ZIKV into Colombia, resulting in varying degrees of sampled onward transmission.

### Multiple introductions of ZIKV to Colombia

We infer patterns of ZIKV introduction and spread in Colombia from the phylogenetic placement of 20 Colombian sequences. Previously, 12 of these sequences were publicly available. We add an additional 8 Colombian sequences sampled over a broad geographic and temporal range. Colombian ZIKV sequences clustered into two distinct clades (Figure 3B). Both clades are descended from viruses inferred to be from Brazil (Figure 3A). Viruses immediately ancestral to both Colombian clades are estimated by the phylogeographic model to have 100% model support for a Brazilian origin. However, lack of genomic sampling from many ZIKV-affected countries in the Americas may limit our ability to infer direct introduction from Brazil or transmission through unsampled countries prior to arrival in Colombia.

Clade 1 is comprised of 28 viruses, 17 of which are from Colombia, and is characterized by nucleotide mutations T738C, C858T, G864T, C3442T, A3894G, C5991T, C9279T, and A10147G. This clade contains all previously reported Colombian genomes, as well as five of the genomes generated during this study (Figures 3 and 4). The phylogeographic model places the root of this clade in Colombia with 99% model support. The most parsimonious reading is that this clade of viruses resulted from a single introduction event from Brazil into Colombia.

**Figure 4.**
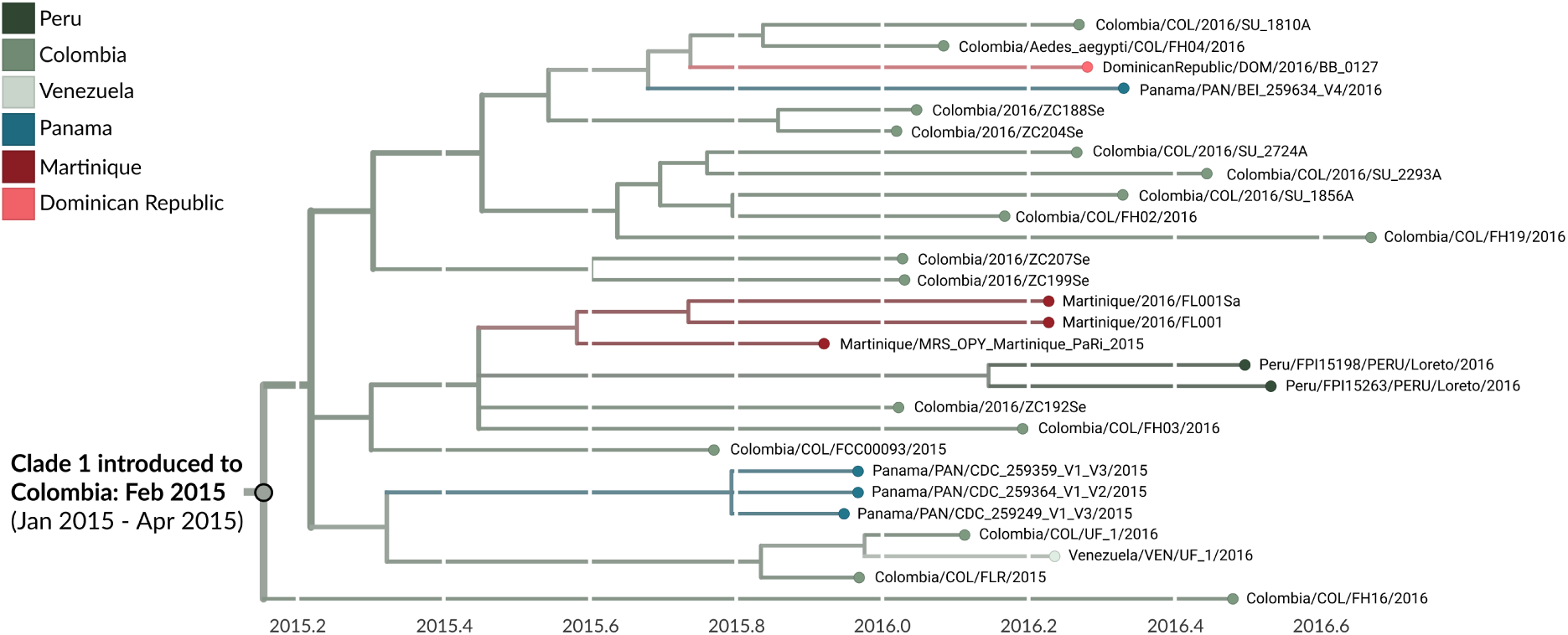
Phylogeny of 28 clade 1 viruses. The inferred geographic migration history indicates movement from Brazil into Colombia, with subsequent migration into other countries in South America, Central America, and the Caribbean. The estimate of introduction timing and the 95% confidence interval are given for the most basal node of the clade.

Clade 2 contains 5 viruses, 3 of which are from Colombia (Figures 3 and 5). All three were sequenced during this study, and thus this clade was not previously recognized in Colombia. This clade is characterized by mutations T1858C, A3780G, G4971T, C5532T, G5751A, A6873G, T8553C, C10098T. The phylogeographic model places the root of this clade in Colombia with 98% model support and also suggests a Brazil to Colombia transmission route that is likely, but not certainly, direct. Two additional genomes with less that 50% genomic coverage also group within these clades; COL/FH14/2016 within clade 1 and COL/FH15/2016 within clade 2 (data not shown).

**Figure 5.**
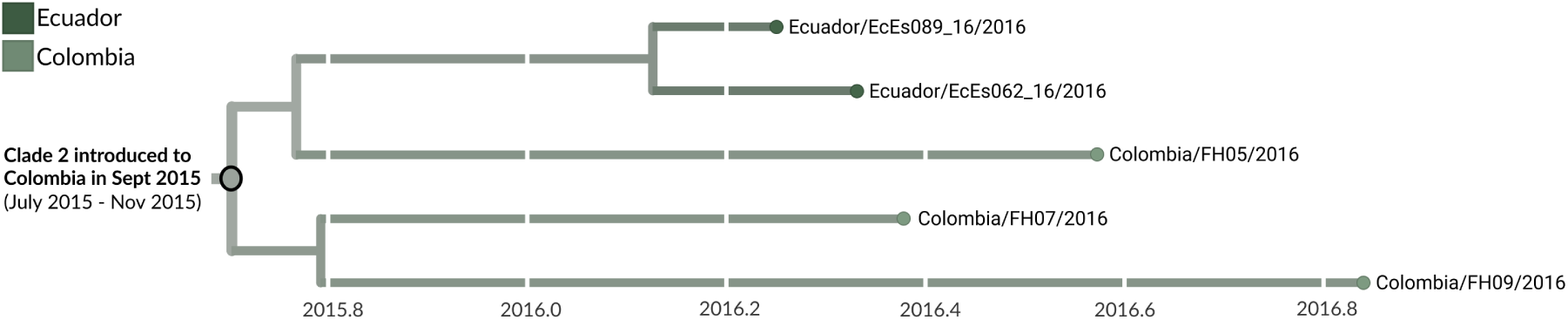
Phylogeny of 5 clade 2 viruses. The inferred geographic migration history indicates movement from Brazil into Colombia, with subsequent migration into Ecuador as evidenced by the nesting of Ecuadorian genomes EcEs062 16 and EcEs089 16. The estimate of introduction timing and the 95% confidence interval are given for the most basal node of the clade.

To examine how our estimate of two introductions to Colombia might be affected by the number of sequences available, we conducted rarefaction analyses. Under the assumption that sequenced viruses are sampled at random, these curves show how much genomic sampling is required to fully sample the circulating viral diversity. Figure 6 shows the number of introductions to Colombia or Mexico inferred under the phylogeographic model as a function of the number of sequences sampled from that country. For ZIKV in Mexico, we see that the rarefaction curve begins to flatten once roughly 25 to 30 Mexican viruses have been sampled (Figure 6A). In contrast, for Colombia we do not observe further ZIKV introductions once we have sampled around 4 genomes (Figure 6B). Thus, while there are only 20 ZIKV whole genomes available from Colombia, we think it is unlikely that we would observe more ZIKV introductions given more sequence data.

**Figure 6.**
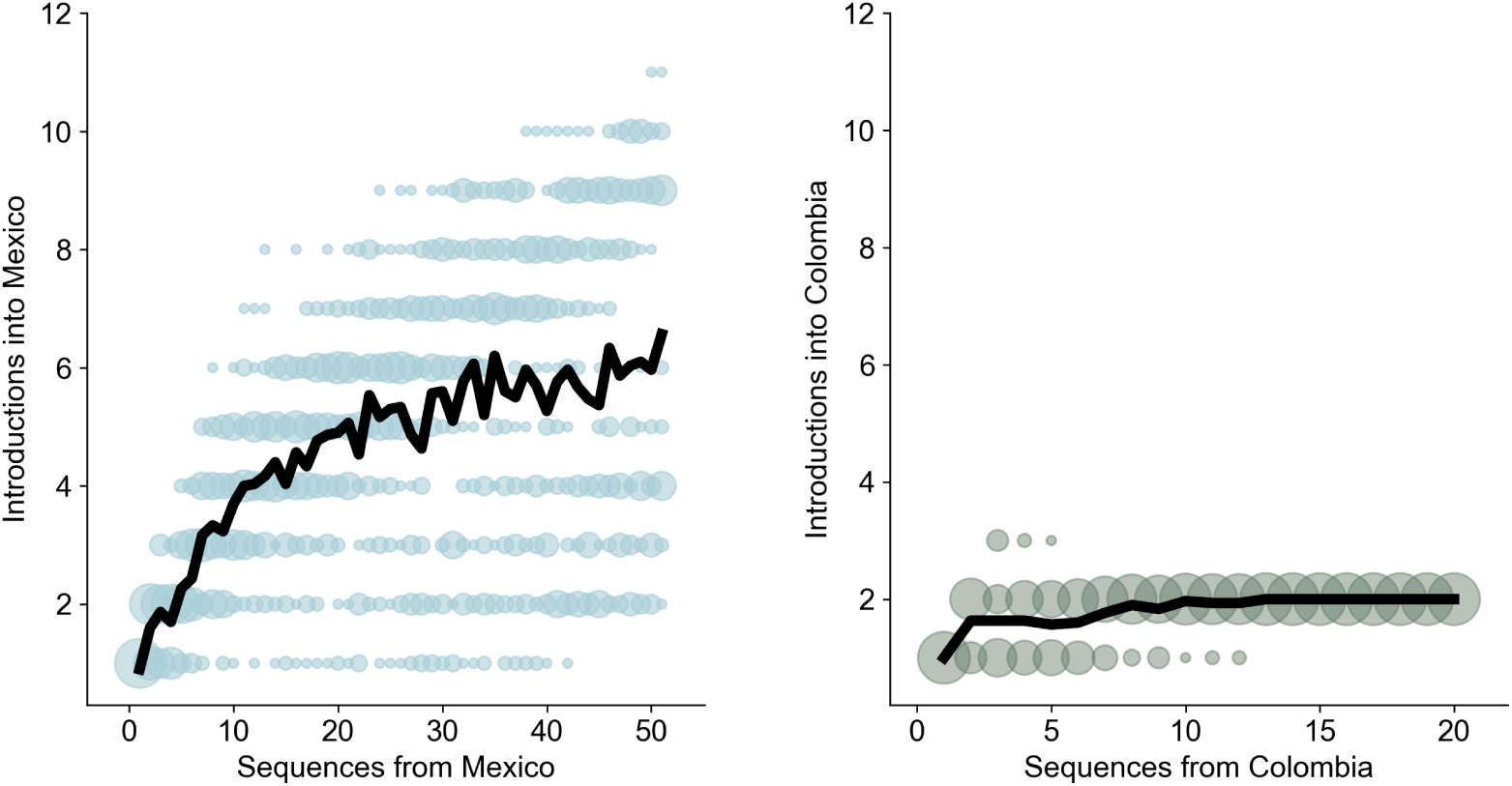
Rarefaction curves for Mexican ZIKV and Colombian ZIKV. For both Mexico and Colombia, the number of introductions into the country is plotted as a function of the number of genomes sampled from that country, where genomes are subsampled from available sequences. The colored circles show introduction counts for all of the phylogeographically labelled trees; they are sized according to the frequency with which a specific number of introductions was observed for a given level of subsampling. The black line shows the mean number of introductions observed for a given level of subsampling.

### Transmission within Colombia

We estimate that clade 1 was introduced to Colombia around late February of 2015 (95% CI: January 2015 to April 2015) (Figures 3B and 4), and that clade 2 was introduced in mid September of 2015 (95% CI: July 2015 to November 2015) (Figures 3B and 5). Our estimate of clade 1 introduction timing supports between five and eight months of cryptic ZIKV transmission within Colombia prior to initial case detection in September 2015, a finding that is consistent with other genomic epidemiological studies of ZIKV [7, 8, 10].

Genomic sampling was available from 7 of 32 departments within Colombia (Figure 7A). Four genomes were sampled from Santander, three viruses were sampled from Valle del Cauca, and two viruses came from Córdoba. One virus each was sampled from Risaralda, Tolima, Huila, and Atlántico departments respectively (Figure 7B and C). Seven of the publicly-available Colombian ZIKV genomes lacked department-level information (Figure 7B). Given that many ZIKV-affected departments within Colombia have only minimal genomic sampling, or lack it all together, we have refrained from using phylogeographic methods to reconstruct the direction of transmission between Colombian departments. However, we do note some signals of geographic influence on transmission within the phylogenies. For example, two closely related clade 1 viruses (COL/2016/SU 1810A and COL/2016/SU 2724A) were both sampled from Santander, and may be linked cases.

**Figure 7.**
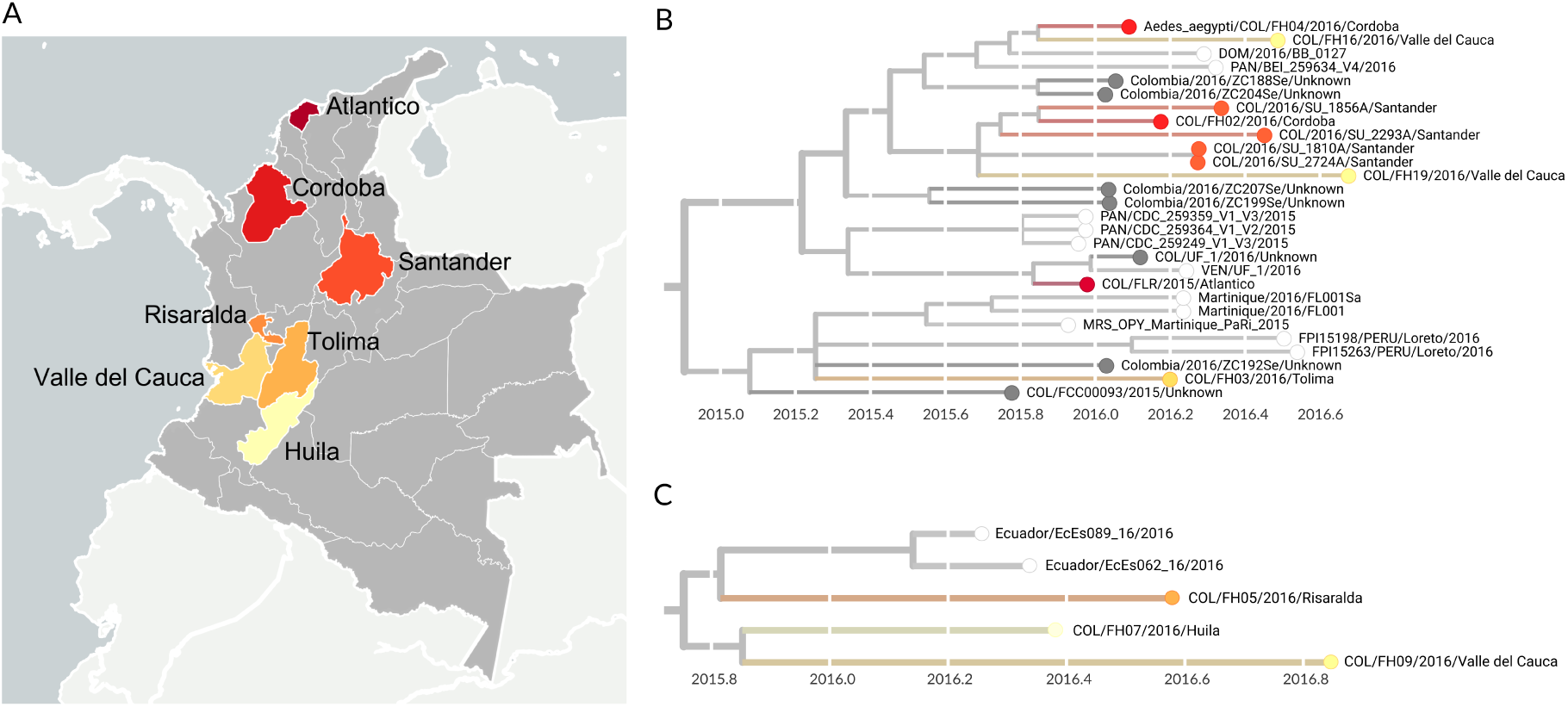
Department-level sampling information for 20 viruses sequenced from Colombia. A) Colombian departments from which Zika virus was sampled are highlighted. These departments included Santander, Valle del Cauca, Córdoba. Risaralda, Tolima, Huila, and Atlántico departments. Seven genomes lack public information about which department they were sampled from. The land-sea mask, coastline, lake, river and political boundary data are extracted from datasets provided by Generic Mapping Tools (GMT) licensed under GNU General Public License. B) Phylogeny of clade 1 viruses, colored by Colombian department of sampling. Viruses not sampled from Colombia are indicated with white tips, and Colombian viruses lacking information about sampling department are colored dark grey. Phylogeny of clade 2 viruses, colored by Colombian department of sampling. Viruses not sampled from Colombia are indicated with white tips.

### Transmission from Colombia to other countries

We find evidence for onward transmission from Colombia into other countries in the Americas. Clade 1 shows movement of viruses into countries that share a border with Colombia, namely Panama, Venezuela, and Peru, as well as into the Dominican Republic and Martinique (Figure 4). Clade 2 indicates movement of Colombian ZIKV into neighboring Ecuador (Figure 5). Transmission from Colombia into bordering countries seems reasonable, and these patterns agree with previously documented trends of ZIKV expansion in the Americas [7, 9, 10], but provide more detail due to the greater amounts of sequence data now available. For instance, analysis by Metsky et al [10] also supports movement of ZIKV from Colombia to Martinique and the Dominican Republic. However, without sequence data from Panama, Peru, or Venezuela, they were unable to capture spread from Colombia into these countries.

## Discussion

Despite the scale of the Colombian epidemic, publicly available sequence data were limited, and no detailed genomic epidemiological analysis of ZIKV dynamics had been performed. We sought to improve genomic sampling for Colombia, and to perform a detailed genomic analysis of the Colombian epidemic. Only 12 Colombian genomes were available prior to this study. To these data we added 8 new sequences sampled broadly across Colombia, and performed a phylogeographic analysis of American ZIKV. We describe general transmission patterns across the Americas and present estimates of ZIKV introduction timing and frequency specific to Colombia. We find evidence of at least two introductions of ZIKV to Colombia, yet remarkably the majority of Colombian viruses cluster within a single clade, indicating that a single introduction event caused the majority of ZIKV cases in Colombia. Under the assumption that viruses sequenced from Colombia are random samples of Colombian ZIKV cases, we find that Colombian ZIKV diversity is well represented by 20 Colombian genomes. It is therefore unlikely that further genomic sampling would reveal more introductions of ZIKV into Colombia. ZIKV dispersal out of Colombia also appears widespread, with movement to bordering countries (Panama, Venezuela, Ecuador, and Peru) as well as more distal countries in the Caribbean.

While it may be tempting to read the inferred phylogeographic migration history as a complete record of transmission between countries, we caution against doing this for analyses of ZIKV. In contrast to other large outbreaks, such as the Ebola epidemic in West Africa, genomic sampling of the American ZIKV epidemic is sparse. Many ZIKV-affected countries have minimal genomic sampling; others have none at all. Thus while the phylogeographic model will correctly infer the geographic location of internal nodes given the dataset at hand, adding sequences from previously unsampled countries may alter migration histories such that apparent direct transmission from country A to country C instead becomes transmission from country A to country B to country C.

Consistent with other studies, our estimates of when these introductions occurred support cryptic ZIKV transmission prior to initial case confirmation. Perhaps more surprisingly, our estimate of the age of clade 1 indicates that ZIKV likely spread to Colombia even before official confirmation of ZIKV circulation in Brazil [37]. These findings underscore the utility of genomic epidemiology to date introduction events and describe transmission patterns that are difficult to detect using traditional surveillance methods, thereby providing more accurate definitions of the population at risk and a better understanding of how importation and within-country transmission shape epidemics.

## List of abbreviations

ZIKV: Zika virus
rRT-PCR: real time reverse-transcription polymerase chain reaction
Ct value: Cycle threshold value

## Declarations

### Ethics approval and consent to participate

In Colombia, most ZIKV diagnostic testing occurred at the INS. However, Colombian academics were also involved in ZIKV surveillance, sampling *Ae.aegypti* and limited human cases. Our study includes samples collected by both the INS and the Universidad del Rosario (UR). Sample collection by UR was approved by the Universidad del Rosario Ethics Committee, approval number 617-2017. Written consent was obtained for human specimen collection. Sample collection performed by the INS was approved by the Comité de Ética y Metodologías de la Investigación (CEMIN) - Instituto Nacional de Salud, approval number CEMIN 39-2017. Consent for specimen collection was waived as specimens were collected as part of the Colombian public health response. All sample selection for inclusion in the genomic analysis, sequencing library preparation, and data analysis were performed on anonymized samples and data.

### Consent for publication

Not applicable.

### Availability of data and material

All priming schemes, kit specifications, laboratory protocols, bioinformatic pipelines, and sequence validation information are openly available at github.com/blab/zika-seq. Nextstrain analytic and visualization code is available at github.com/nextstrain. Code and builds specific to the analysis presented here, as well as all genomes sequenced for this study, are openly available at github.com/blab/zika-colombia. Consensus sequences are also available on NCBI GenBank (accessions MK049245 through MK049252). The phylogeny and all inferences reported here can be interactively explored at nextstrain.org/community/blab/zika-colombia.

## Authors’ contributions

AB, LHM, DPR, IML, MEH, JDR, MMR, and TB conceived of the study and contributed to study design. AB, LHM, KLD, LP, AR, and CT, performed the lab work. AB and BP performed the bioinformatic assembly. AB performed the phylogeographic analysis. AB, LHM, and TB interpreted the data and wrote the manuscript. DPR, MEH, DPC, JDR, and MMR facilitated the collaboration and sought ethics approvals for the work. JDR, DPC, and MMR provided clinical samples and metadata. All authors provided critical feedback that helped shape the research and the manuscript.

## Acknowledgements

We thank R. Shabman, B. Pickett, P. Rahal, L. Karan, R. Delgado, A. Enfissi, N. Grubaugh, and R. Lanciotti for giving us permission to include unpublished genomes available on GenBank in our analysis. We also thank Maria Fernanda Torres for facilitating this collaboration and Adam Geballe for generously loaning space in his laboratory at the Fred Hutch.

## References

1. PAHO. Zika Cumulative Cases;. https://www.paho.org/hq/index.php?option=com_content&view=article&id=12390:zika-cumulative-cases&Itemid=42090&lang=en.

2. Duffy MR, Chen TH, Thane Hancock W, Powers AM, Kool JL, Lanciotti RS, et al. Zika Virus Outbreak on Yap Island, Federated States of Micronesia. N Engl J Med. 2009;360(24):2536–2543.

3. Zambrana JV, Bustos Carrillo F, Burger-Calderon R, Collado D, Sanchez N, Ojeda S, et al. Seroprevalence, risk factor, and spatial analyses of Zika virus infection after the 2016 epidemic in Managua, Nicaragua. Proc Natl Acad Sci U S A. 2018;115(37):9294–9299.

4. Aubry M, Teissier A, Huart M, Merceron S, Vanhomwegen J, Roche C, et al. Zika Virus Seroprevalence, French Polynesia, 2014–2015. Emerg Infect Dis. 2017;23(4):669–672.

5. Netto EM, Moreira-Soto A, Pedroso C, Höser C, Funk S, Kucharski AJ, et al. High Zika Virus Seroprevalence in Salvador, Northeastern Brazil Limits the Potential for Further Outbreaks. MBio. 2017;8(6).

6. Lessler J, Chaisson LH, Kucirka LM, Bi Q, Grantz K, Salje H, et al. Assessing the global threat from Zika virus. Science. 2016;353(6300):aaf8160.

7. Faria NR, Quick J, Claro IM, Thézé J, de Jesus JG, Giovanetti M, et al. Establishment and cryptic transmission of Zika virus in Brazil and the Americas. Nature. 2017;546(7658):406–410.

8. Grubaugh ND, Ladner JT, Kraemer MUG, Dudas G, Tan AL, Gangavarapu K, et al. Genomic epidemiology reveals multiple introductions of Zika virus into the United States. Nature. 2017;546(7658):401–405.

9. Thézé J, Li T, du Plessis L, Bouquet J, Kraemer MUG, Somasekar S, et al. Genomic Epidemiology Reconstructs the Introduction and Spread of Zika Virus in Central America and Mexico. Cell Host Microbe. 2018;23(6):855–864.e7.

10. Metsky HC, Matranga CB, Wohl S, Schaffner SF, Freije CA, Winnicki SM, et al. Zika virus evolution and spread in the Americas. Nature. 2017;546(7658):411–415.

11. Faria NR, Azevedo RdSdS, Kraemer MUG, Souza R, Cunha MS, Hill SC, et al. Zika virus in the Americas: Early epidemiological and genetic findings. Science. 2016;352(6283):345–349.

12. Calvet G, Aguiar RS, Melo ASO, Sampaio SA, de Filippis I, Fabri A, et al. Detection and sequencing of Zika virus from amniotic fluid of fetuses with microcephaly in Brazil: a case study. Lancet Infect Dis. 2016;16(6):653–660.

13. Naccache SN, Thézé J, Sardi SI, Somasekar S, Greninger AL, Bandeira AC, et al. Distinct Zika Virus Lineage in Salvador, Bahia, Brazil. Emerg Infect Dis. 2016;22(10):1788–1792.

14. Lednicky J, Beau De Rochars VM, El Badry M, Loeb J, Telisma T, Chavannes S, et al. Zika Virus Outbreak in Haiti in 2014: Molecular and Clinical Data. PLoS Negl Trop Dis. 2016;10(4):e0004687.

15. Lanciotti RS, Lambert AJ, Holodniy M, Saavedra S, Signor LDCC. Phylogeny of Zika Virus in Western Hemisphere, 2015. Emerg Infect Dis. 2016;22(5):933–935.

16. Enfissi A, Codrington J, Roosblad J, Kazanji M, Rousset D. Zika virus genome from the Americas. Lancet. 2016;387(10015):227–228.

17. Guerbois M, Fernandez-Salas I, Azar SR, Danis-Lozano R, Alpuche-Aranda CM, Leal G, et al. Outbreak of Zika Virus Infection, Chiapas State, Mexico, 2015, and First Confirmed Transmission by Aedes aegypti Mosquitoes in the Americas. J Infect Dis. 2016;214(9):1349–1356.

18. Pessôa R, Patriota JV, Lourdes de Souza Md, Felix AC, Mamede N, Sanabani SS. Investigation Into an Outbreak of Dengue-like Illness in Pernambuco, Brazil, Revealed a Cocirculation of Zika, Chikungunya, and Dengue Virus Type 1. Medicine. 2016;95(12):e3201.

19. Giovanetti M, Milano T, Alcantara LC, Carcangiu L, Cella E, Lai A, et al. Zika Virus spreading in South America: Evolutionary analysis of emerging neutralizing resistant Phe279Ser strains. Asian Pac J Trop Med. 2016;9(5):445–452.

20. Quick J, Grubaugh ND, Pullan ST, Claro IM, Smith AD, Gangavarapu K, et al. Multiplex PCR method for MinION and Illumina sequencing of Zika and other virus genomes directly from clinical samples. Nat Protoc. 2017;12(6):1261–1276.

21. Pacheco O, Beltrán M, Nelson CA, Valencia D, Tolosa N, Farr SL, et al. Zika Virus Disease in Colombia - Preliminary Report. N Engl J Med. 2016;.

22. Rojas DP, Dean NE, Yang Y, Kenah E, Quintero J, Tomasi S, et al. The epidemiology and transmissibility of Zika virus in Girardot and San Andres island, Colombia, September 2015 to January 2016. Euro Surveill. 2016;21(28).

23. Instituto Nacional de Salud. Boletin epidemiologico semanal - semana epidemiologica 08 de 2016;. https://www.ins.gov.co/buscador-eventos/BoletinEpidemiologico/2016%20Bolet%C3%ADn%20epidemiol%C3%B3gico%20semana%208.pdf.

24. Cuevas EL, Tong VT, Rozo N, Valencia D, Pacheco O, Gilboa SM, et al. Preliminary Report of Microcephaly Potentially Associated with Zika Virus Infection During Pregnancy - Colombia, January-November 2016. MMWR Morb Mortal Wkly Rep. 2016;65(49):1409–1413.

25. Moore CA, Staples JE, Dobyns WB, Pessoa A, Ventura CV, Fonseca EBd, et al. Characterizing the Pattern of Anomalies in Congenital Zika Syndrome for Pediatric Clinicians. JAMA Pediatr. 2017;171(3):288–295.

26. Parra B, Lizarazo J, Jiménez-Arango JA, Zea-Vera AF, González-Manrique G, Vargas J, et al. Guillain-Barré Syndrome Associated with Zika Virus Infection in Colombia. N Engl J Med. 2016;375(16):1513–1523.

27. Santiago GA, Vázquez J, Courtney S, Matías KY, Andersen LE, Colón C, et al. Performance of the Trioplex real-time RT-PCR assay for detection of Zika, dengue, and chikungunya viruses. Nat Commun. 2018;9(1).

28. Faye O, Faye O, Diallo D, Diallo M, Weidmann M, Sall AA. Quantitative real-time PCR detection of Zika virus and evaluation with field-caught mosquitoes. Virol J. 2013;10:311.

29. Li H, Durbin R. Fast and accurate short read alignment with Burrows-Wheeler transform. Bioinformatics. 2009;25(14):1754–1760.

30. Pickett BE, Sadat EL, Zhang Y, Noronha JM, Burke Squires R, Hunt V, et al. ViPR: an open bioinformatics database and analysis resource for virology research. Nucleic Acids Res. 2011;40(D1):D593–D598.

31. Hadfield J, Megill C, Bell SM, Huddleston J, Potter B, Callender C, et al. Nextstrain: real-time tracking of pathogen evolution. Bioinformatics. 2018;.

32. Katoh K, Misawa K, Kuma KI, Miyata T. MAFFT: a novel method for rapid multiple sequence alignment based on fast Fourier transform. Nucleic Acids Res. 2002;30(14):3059–3066.

33. Nguyen LT, Schmidt HA, von Haeseler A, Minh BQ. IQ-TREE: a fast and effective stochastic algorithm for estimating maximum-likelihood phylogenies. Mol Biol Evol. 2015;32(1):268–274.

34. Sagulenko P, Puller V, Neher RA. TreeTime: Maximum-likelihood phylodynamic analysis. Virus Evol. 2018;4(1):vex042.

35. Suchard MA, Lemey P, Baele G, Ayres DL, Drummond AJ, Rambaut A. Bayesian phylogenetic and phylodynamic data integration using BEAST 1.10. Virus Evol. 2018;4(1):vey016.

36. Pettersson JHO, O Pettersson JH, Eldholm V, Seligman SJ, Lundkvist Å, Falconar AK, et al. How Did Zika Virus Emerge in the Pacific Islands and Latin America? MBio. 2016;7(5).

37. PAHO. Epidemiological Alert: Zika virus infection. PAHO; 2015.

